# White matter alterations in glaucoma and vision-deprived brains differ outside the visual system

**DOI:** 10.1101/2020.11.30.404434

**Authors:** Sandra Hanekamp, Branislava Ćurčić-Blake, Bradley Caron, Brent McPherson, Anneleen Timmer, Doety Prins, Christine C. Boucard, Masaki Yoshida, Masahiro Ida, Nomdo M. Jansonius, Franco Pestilli, Frans W. Cornelissen

## Abstract

The degree to which glaucoma has effects beyond the eye –in the brain– is unclear. We investigated white matter microstructure (WMM) alterations in 37 tracts of patients with glaucoma, monocular blindness and controls. We used reproducible methods and the advanced cloud computing platform brainlife.io. White matter tracts were subdivided into seven categories ranging from primarily involved in vision (the visual white matter) to primarily involved in cognition and motor control. WMM in both glaucoma and monocular blind subjects was lower than controls in the visual white matter, suggesting neurodegenerative mechanisms due to reduced sensory inputs. In glaucoma participants WMM differences from controls decreased outside the visual white matter. A test-retest validation approach was used to validate these results. The pattern of results was different in monocular blind participants, where WMM properties increased outside the visual white matter as compared to controls. The pattern of results suggests that whereas in the blind loss of visual input might promote white matter reorganization outside of the early visual system, such reorganization might be reduced or absent in glaucoma. The results provide indirect evidence that in glaucoma unknown factors might limit the brain plasticity effects that in other patient groups follow visual loss.

## Introduction

Glaucoma is a chronic and progressive disease characterized by damage to the retinal nerve fiber layer (RNFL) and associated with morphological changes to the optic nerve(“European Glaucoma Society Terminology and Guidelines for Glaucoma, 4th Edition - Chapter 3: Treatment Principles and Options Supported by the EGS Foundation: Part 1: Foreword; Introduction; Glossary; Chapter 3 Treatment Principles and Options,” 2017). The traditional view is that high levels of intraocular pressure (IOP) are not tolerated by the retina and optic nerve resulting in a deformed lamina cribrosa, causing axonal damage and consequent apoptotic death of retinal ganglion cells (RGCs). However, over the last decade, research has shown that glaucoma might not be solely an eye disease, but a complex neurodegenerative disease also involving processes within the brain tissue (Bagetta & Nucci, 2015; Danesh-Meyer & Levin, 2015; Walsh, 2020). While the nature of the neurodegenerative component to glaucoma remains to be resolved, it is clear that damage in glaucoma is not restricted to the axons that form the optic nerve. In addition, numerous magnetic resonance imaging (MRI) studies have shown that the pre-geniculate (optic tract and chiasm), geniculate, and post-geniculate (the optic radiation (OR) and primary visual cortex) structures are affected as well in glaucoma, at least in later stages of the disease (C. Boucard et al., 2016; C. C. Boucard et al., 2009; W. W. Chen et al., 2013; Z. Chen et al., 2013; Dai et al., 2013; Garaci et al., 2009; Haykal et al., 2019; Hernowo et al., 2011; C. Li et al., 2012; K. Li et al., 2014; Liu et al., 2012; Murai et al., 2013; Prins et al., 2016; M.-Y. Wang et al., 2013; Yu et al., 2013; Zikou et al., 2012). Moreover, changes in white matter (WM) tracts beyond the primary visual pathways such as the anterior thalamic radiation, corticospinal tract, superior longitudinal fasciculus, and forceps major (C. Boucard et al., 2016; Frezzotti et al., 2014; Trivedi et al., 2019; J. Wang et al., 2016; R. Wang et al., 2018; Zikou et al., 2012) and a network reorganization of the glaucomatous brain (Di Ciò et al., 2020). In line with this idea, it has also been proposed that glaucoma and Alzheimer’s Disease (AD) might share a common pathogenesis (Mancino et al., 2018). Indeed, both diseases are progressive, chronic, age-related diseases and cause RGC degeneration and irreversible neuronal cell loss. Furthermore, research suggests a genetic link between glaucoma and AD (Cumurcu et al., 2013; Inoue et al., 2013; Tamura et al., 2006). More recently, dysfunction of the glymphatic system (i.e. a cerebrospinal fluid transport system that facilitates clearance of neurotoxic molecules) was proposed as a novel and missing link between AD and glaucoma (Mathieu et al., 2018; Sen et al., 2020; X. Wang et al., 2020; Wostyn, 2020; Wostyn et al., 2015, 2017).

At the present, the most parsimonious explanation for the changes to the brain tissue in glaucoma remains that they are a consequence of the decreased visual input due to retinal and optic nerve damage (Nauta & Ebbesson, 1970; Prins et al., 2016). Yet, a complete picture of the alterations of the WM tissue in glaucoma is missing. Understanding the degree to which the alterations in the brain tissue in glaucoma go beyond the early visual system, and the degree to which the effect of glaucoma outside of the visual system is similar to that of other diseases also affecting visual input can help to clarify the pathophysiology of glaucoma. Answering this question is critical, to advance our understanding of the disease and may, ultimately, inform future therapeutic approaches.

The current study tested whether the WM alterations are unique to the glaucomatous brain or depend on a decreased visual input. To do so, the study used reproducible cloud computing methods to measure WM microstructure in a larger set of tracts (Avesani et al., 2019; Bullock et al., 2019) than previously done. We compared WM microstructure between two independent sets of glaucoma patients (using the independence and demographic differences in the two glaucoma data sets to replicate the pattern of results) and one monocular blind patients group. We note that blindness was defined as: “Light-perception negative for at least five years, due to perforation, enucleation, evisceration or ablatio retinae leading to blindness to the affected eye”. For the purpose of our study, monocular blindness acquired later in life was considered a reference model for the effect of reduced visual input to the brain. This is because due to unilateral sensory deafferentation half of the normal visual input is lost in these patients (Nys et al., 2015).

## Results

The goal of this study was twofold. First, we wanted to determine the extent of the WM alterations in the glaucomatous brain outside of the visual WM (Rokem et al., 2017). Second, we wanted to establish whether the pattern of alterations across a large set of WM tracts was unique to glaucoma or was instead similar to those observed in a model of reduced visual input (i.e monocular blindness). A total of 102 subjects, including subjects with glaucoma (GL), monocular blindness (MBL) and healthy controls (HC). To validate our results in glaucoma, we used two independent data sets with major differences in the demographics of the participants (one group from the Netherlands and another from Japan, see **Methods: subjects**). This resulted in a total of two glaucoma patients groups (GL1 and GL2) and three control groups (MBL, HC1 and HC2). The demographics and clinical characteristics are summarized for the first cohort in **Table 1** and the second cohort in **Table 2** in the **Methods** section. The participant groups did not differ in mean age (GL1 versus HC1: p=0.80, MBL versus HC1: p=0.65, GL2 versus HC2: p=0.90). The gender distribution did not differ in the NL participants (GL1 versus HC1: p=0.52, MBL versus HC1: p=0.73), but differed in the JP participants (GL2 and HC2 p=0.04).

**Table 1.** Characteristics of Cohort 1 (collection site at Netherlands; NL)

|  | **HC1 (N=20)** | **GL1 (N=17)** | **MBL (N=14)** |
| --- | --- | --- | --- |
|  | Mean (Min/Max)<br>or percentage | Mean (Min/Max)<br>or percentage | Mean (Min/Max)<br>or percentage |
| Age | 64.05 (53.0 to 81) | 63.29 (53 to 82) | 63(54 to 72) |
| Male gender, % | 60% | 47.06% | 50% |
| Duration defect, yrs |  | 6.47 (1 to 14) | 18.79 (6 to 60) |
| **VFMD, dB** |  |  |  |
| Better eye |  | -1.52 (-4.81 to 0.46) |  |
| Worse eye |  | -20.33 (-30.6 to -6.56) |  |
| Mean eye |  | -10.93 (-16.81 to -3.4) |  |
| **NFI** |  |  |  |
| Better eye | 17.35 (6 to 29) | 25.86 (3 to 64) | 19.43 (2 to 58) |
| Worse eye | 27.5 (7 to 51) | 75.64 (28 to 98) |  |
| Mean eye | 22.43 (6.5 to 36) | 50.75 (26 to 81) |  |
| **IOP, mmHg** |  |  |  |
| Better eye | 14.53 (10 to 20) | 14.71 (10 to 20) | 15.5 (12 to 21) |
| Worse eye | 15 (10 to 25) | 13.57 (4 to 18) |  |
| Mean eye | 14.9 (10 to 22.5) | 13.41 (6.5 to 18.5) |  |
| VFMD= mean deviation (dB), NFI= nerve fiber indicator, IOP= intraocular pressure |  |  |  |

**Table 2.**
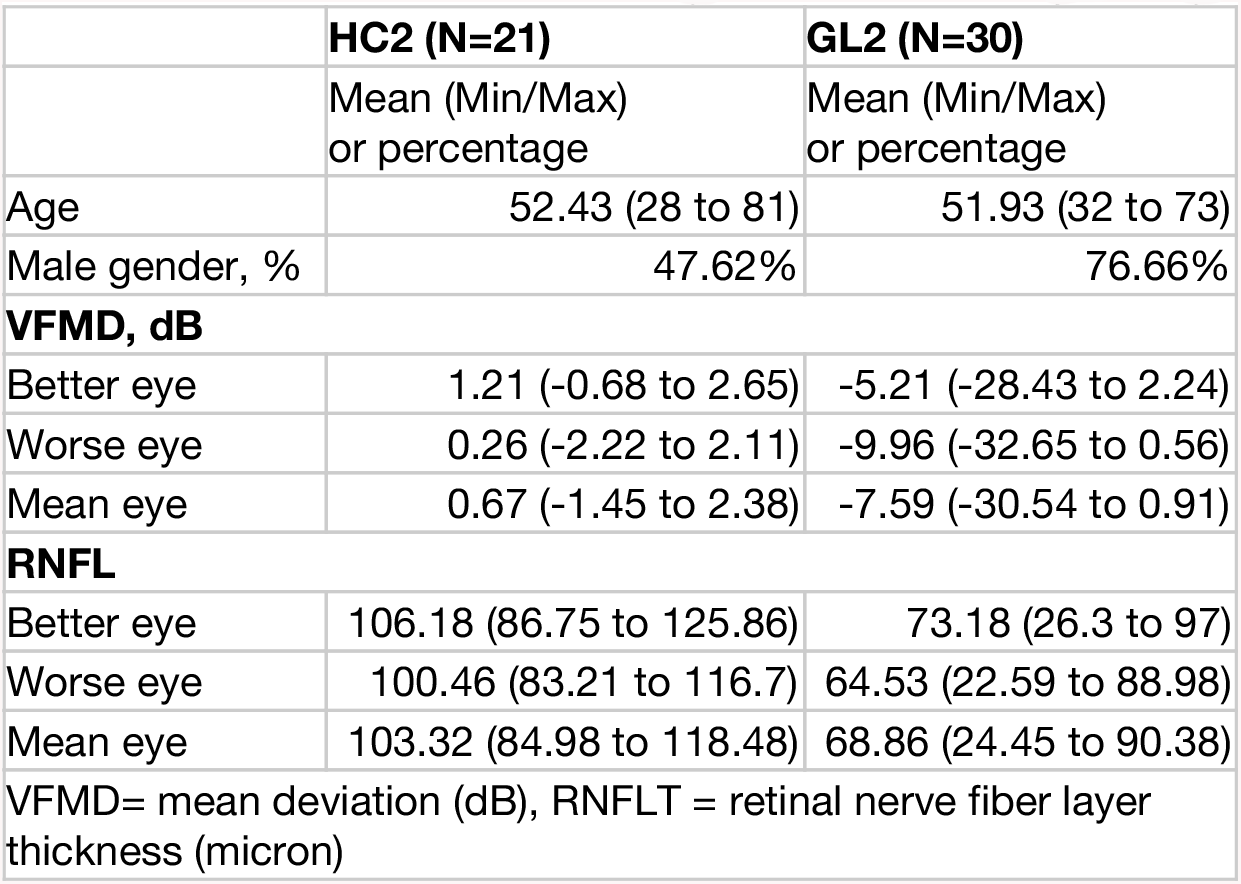
Characteristics of Cohort 2 (collection site at Japan; JP)

Diffusion-weighted magnetic resonance imaging data was processed using an automated pipeline implemented as reproducible web-services on brainlife.io (see **Table 3** for the full processing pipeline (Ades-Aron et al., 2018; Avesani et al., 2019; McPherson & Pestilli, n.d.)). Using this pipeline, thirty seven major WM tracts were identified (**Figure 1A** right-hand side; see also **Table 4**) (Bullock et al., 2019; Wassermann et al., 2016). After segmentation of all tracts, we computed WM microstructure (WMM) fractional anisotropy (FA) and mean diffusivity (MD) along the tract-length (**Figure 1A** left-hand side; see also **Methods: Tract profile generation**) (Peter J. Basser & Pierpaoli, 2011; Yeatman et al., 2012).

**Table 3.** Description and web-links to the open source code and open cloud services used in the creation of this dataset.

| Application | Source Code | Open Cloud Service |
| --- | --- | --- |
| **Anatomy** |  |  |
| FSL Anat (T1) | github.com/brainlife/app-fsl-anat/tree/v1.0 | doi.org/10.25663/brainlife.app.273 |
| Freesurfer | github.com/brainlife/app-freesurfer/tree/1.8 | doi.org/10.25663/bl.app.0 |
| Tissue-type segmentation | github.com/brainlife/app-mrtrix3-5tt | doi.org/10.25663/brainlife.app.239 |
| Generate ROIs for Visual White Matter Tracking in dMRI Space (Benson pRF) | github.com/brainlife/app-roigenerator/tree/optic-radiation-v1.0 | doi.org/10.25663/brainlife.app.349 |
| pRFs/ Benson14-Retinotopy | github.com/davhunt/app-benson14-retinotopy/tree/master | doi.org/10.25663/brainlife.app.187 |
| **Diffusion** |  |  |
| Mrtrix3 preprocess | github.com/brainlife/app-mrtrix3-preproc/tree/1.6 | doi.org/10.25663/bl.app.68 |
| Fit CSD Model For Tracking | github.com/bacaron/app-mrtrix3-act/tree/csd_generation-v1.0 | doi.org/10.25663/brainlife.app.238 |
| ACT Tractography | github.com/brainlife/app-mrtrix3-act/tree/1.4 | doi.org/10.25663/brainlife.app.297 |
| White Matter Anatomy Segmentation | github.com/brainlife/app-wmaSeq/tree/3.6 | doi.org/10.25663/brainlife.app.188 |
| Track the Human Optic Radiation (THORA) - MRtrix3 iFOD2 (dwi) | github.com/brainlife/app-mrtrix3-roi2roi-tracking/tree/thora-v1.0 | doi.org/10.25663/brainlife.app.347 |
| Remove Tract Outliers | github.com/brainlife/app-removeTractOutliers/tree/1.3 | doi.org/10.25663/brainlife.app.195 |
| Tract Analysis Profiles | github.com/brainlife/app-tractanalysisprofiles/tree/1.8 | doi.org/10.25663/brainlife.app.361 |
| **QA** |  |  |
| Generate images of T1 | github.com/brainlife/app-slicer-fsl/tree/master-app-v1.0.0 | doi.org/10.25663/brainlife.app.300 |
| Generate images of tissue type masks | github.com/brainlife/app-slicer-fsl/tree/5tt-v1.0.0 | doi.org/10.25663/brainlife.app.312 |
| Generate images of DWI | github.com/brainlife/app-slicer-fsl/tree/master-app-v1.0.0 | doi.org/10.25663/brainlife.app.298 |
| Generate images of DWI overlaid on T1 | github.com/brainlife/app-slicer-fsl/tree/dwi-t1-v1.0.0 | doi.org/10.25663/brainlife.app.309 |
| Plot response function | github.com/brainlife/app-plot-response/tree/v1.0 | doi.org/10.25663/brainlife.app.311 |
| Generate images of tensor | github.com/brainlife/app-slicer-fsl/tree/tensor-v1.0.0 | doi.org/10.25663/brainlife.app.302 |
| Compute SNR on Corpus Callosum | github.com/davhunt/app-snr_in_cc/tree/plot | doi.org/10.25663/brainlife.app.120 |
| Tractography quality check | github.com/brainlife/app-tractographyQualityCheck/tree/1.2 | doi.org/10.25663/brainlife.app.189 |

**Table 4.** White matter tracts of interest and their classification. White matter tracts were categorized based on their involvement in vision. Categorisations were made as follows: 1) early vision tracts are part of the visual pathways, 2) ventral stream tracts are associated with ventral stream processing, 3) dorsal stream tracts are associated with dorsal stream processing, 4) occipital tracts are defined as originating from or projecting to the occipital lobe, 5) vertical tracts are defined as vertically-oriented tracts, and 6) callosal tracts are projecting through the body or genu of the corpus callosum 7) other tracts are defined as WM tracts that are not categorized in previous categories.

| Color | Tract N=(37) | Category |
| --- | --- | --- |
|  | L/R Optic Radiation | Early Vision |
|  | L/R Inferior Frontal Occipital Fasciculus | Ventral Stream |
|  | L/R Inferior Longitudinal Fasciculus | Ventral Stream |
|  | L/R Ventral Occipital Fasciculus | Occipital |
|  | Corpus Callosum - Forceps Major | Occipital |
|  | L/R Superior Longitudinal Fasciculus 1 And 2 | Dorsal Stream |
|  | L/R Superior Longitudinal Fasciculus 3 | Dorsal Stream |
|  | L/R Cingulum Cingulate | Dorsal Stream |
|  | L/R Medial Longitudinal Fasciculus - Angular Gyrus | Vertical |
|  | L/R Medial Longitudinal Fasciculus - Superior Parietal Lobule | Vertical |
|  | L/R P Arcuate Fasciculus | Vertical |
|  | L/R Temporo-parietal Connection | Vertical |
|  | Corpus Callosum - Forceps Minor | Callosal (Non-visual) |
|  | Corpus Callosum - Anterior Frontal Area | Callosal (Cognitive) |
|  | Corpus Callosum - Middle Frontal Area | Callosal (Motor) |
|  | Corpus Callosum - Parietal Area | Callosal (Associative) |
|  | L/R Arcuate Fasciculus | Other |
|  | L/R Frontal Aslant Tract | Other |
|  | L/R Corticospinal Tract | Other |
|  | L/R Fronto Thalamic | Other |
|  | L/R Uncinate Fasciculus | Other |

**Figure 1.**
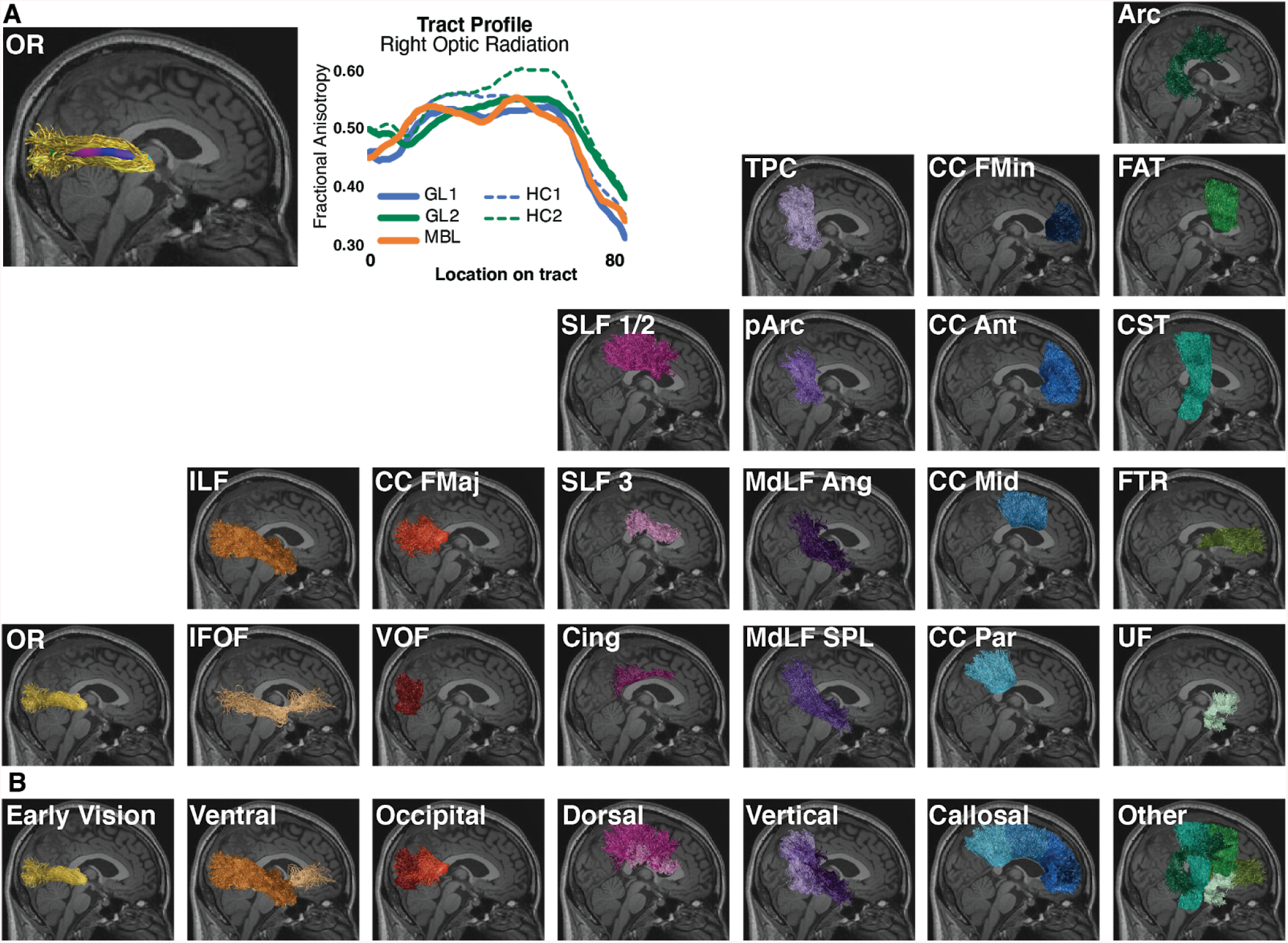
White matter anatomy segmentation. **A.** *Left-hand side*. The 37 WM tracts identified using the brainlife.io white matter segmentation (brainlife.app.188). *Right-hand side*. One representative subject white matter tract (the optic radiation) with overlayed microstructural estimate and representative optic radiation average tract profile for each participant group. Tract profiles were estimated using brainlife.io (brainlife.app.361; (Yeatman et al., 2012)). **B.** *White matter tracts categories*. Tracts were subdivided into 7 categories. These categories were defined by the degree of involvement in vision. Only WM tracts from the right hemisphere are visualized here for a representative subject.

The 37 WM tracts were also assigned to 7 categories varying from tracts highly related to vision and tract more clearly related to cognition and motor control (**Figure 1B**; see also **Table 4**). Categories were defined as follows: (1) *Early vision:* the optic radiation, OR (Rokem et al., 2017), (2) *Ventral visual stream*: Inferior Lateral Fasciculus (ILF) and Inferior Fronto Occipital Fasciculus (IFOF) (Goodale & David Milner, 1992), (3) *Occipital vertical tracts* (associative tracts within the occipital lobe): The vertical occipital fasciculus (VOF) and Forceps Major (CC FMaj) (Takemura et al., 2017), (4) *Dorsal stream tracts*: Superior Lateral Fasciculus (SLF) 1, 2 and 3 and the Cingulum dorsalis (Cing) (Goodale & David Milner, 1992), (5) *Posterior vertical association tracts*: Temporal Parietal Connection (TPC), Posterior Arcuate (pArc), the Middle Longitudinal Fasciculus projecting to the Angular gyrus (MdLF Ang) and projecting to the Superior Parietal Lobule (MdLF SPL) (Bullock et al., 2019), (6) *Cross callosum tracts* (projecting through the body or genu of the corpus callosum): CC Forceps Minor (CC FMin), Anterior (CC Ant), Middle (CC Mid), and parietal (CC Par) (Fabri et al., 2014) and (7) *Other tracts*: Fronto-Thalamic Radiation (FTR), Uncinate Fasciculus (UF), Cortico-Spinal Tract (CST), Frontal Aslant Tract (FAT) and the Arcuate Fasciculus (Arc).

The average FA and MD for each tract in each participant was estimated and used to compute Hedges’ g effect size (independently for FA and MD) (Hentschke & Stüttgen, 2011). To compute g the following participant groups were compared: GL1 and HC1, MBL and HC1 and GL2 and HC2. The average (and standard error) effect size across all 37 WM tracts is reported in **Table S1**(see also **Methods: Statistical analysis**). This procedure returned three sets of 37 estimated *g*. The variable group consisted of three different levels (GL1 vs HC1, MBL vs HC1 and GL2 vs HC2) and the variable category consisted of seven levels (early vision, ventral, occipital, dorsal, vertical, callosal and other tracts (**Figure 1** and **Table 4**). For all the subsequent analyses Two-Way Factorial ANOVAs were conducted independently for FA and MD. The ANOVA compared the main effects of the participants’ group comparisons (see above) and tract category and the interactions between group comparisons and tract category using the Hedges’ g.

### Which white matter tracts show the strongest difference between group comparisons?

To gain insight into which WM tracts were most different in our individual group comparisons, and before discussing the specific statistical tests, we provide a visualization of the Hedges’ g, effect sizes, for all tracts ranked by magnitude (**Figure 2**). We further describe the strongest effect sizes (g>0.7/g<-0.7) based on FA and MD (see also **Table S2**). For the first glaucoma cohort (**Figure 2A**), we found large effect sizes were observed for the right ventral occipital fasciculus (VOF; g=−0.90), bilateral inferior longitudinal fasciculus (ILF; L g= −0.72; R g=−0.72) and bilateral OR (L g= −0.72; R g=−0.70). When comparing MBL vs HC1, the left inferior frontal occipital tract (IFOF; g=0.77) revealed a large effect size. For the second glaucoma cohort, we found large effect sizes for the left OR (g=−0.91), bilateral ILF (L g= −0.76; R g=−0.89) and bilateral (IFOF; L g= −0.82 R g=−0.81). A similar analysis was performed for MD to examine effect sizes for each tract (**Figure 2B**). For our first glaucoma cohort, MD did not reveal large effect sizes. On the other hand, when comparing MBL vs HC1, we found a large effect for the left IFOF (g=−0.83) and left corticospinal tract (g=−0.71). Our second glaucoma cohort revealed large effect sizes for MD in the parietal corpus callosum (g= 0.75) and right OR (g=0.73). As can be seen in **Figure 2A**, when examining FA g estimates, tracts involved in visual processing (e.g OR, ILF, VOF, IFOF) were negatively affected in all group comparisons suggesting decreased FA compared to controls. For monocular blindness, tracts less involved in vision, however, revealed a positive effect size indicating increased FA in comparison to controls. This pattern is uniquely different from glaucoma group comparisons: both glaucoma groups reveal a negative effect on the WMM for the majority of WM tracts, indicating FA is decreased in glaucoma compared to controls.

**Figure 2.**
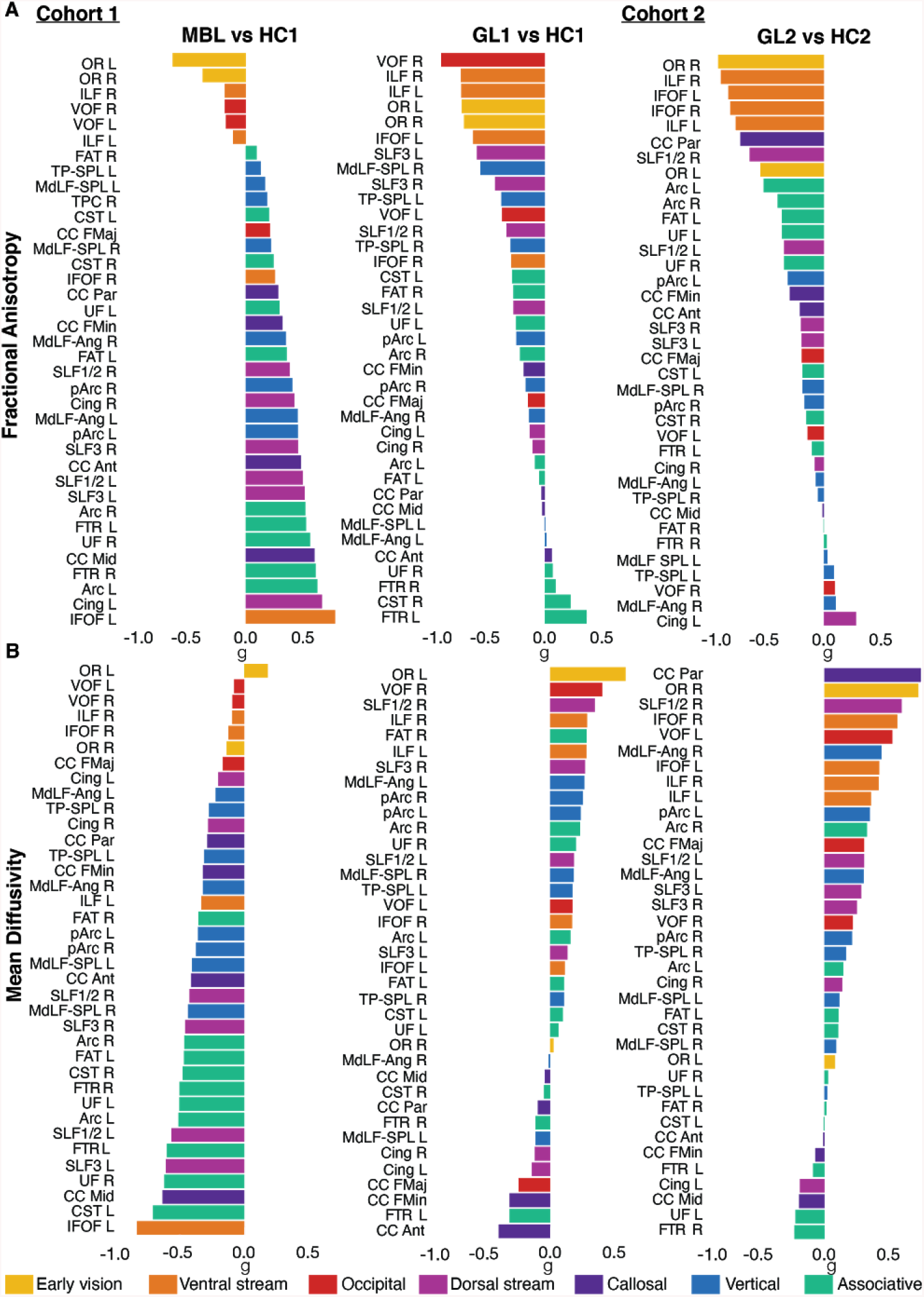
Ordered effect size in glaucoma and monocular blindness for each white matter tract. **A.** Fractional Anisotropy (FA). **B.** Mean Diffusivity (MD). Effect sizes were calculated using Hedges’ g by comparing the average FA or MD values from each WM tract profile. For each white matter tract, FA (or MD) in the glaucoma patients of the first (GL1) or second (GL2) cohort was compared to the corresponding tract in healthy controls (HC1 or HC2 respectively.) Similarly, FA (or MD) in the monocular blind patients (MBL) was compared to the corresponding healthy controls (HC1). Tracts are ordered by effect size with positive effect size presented first (top). WM tract acronyms are fully described in the **Supplementary Information**.

For MD g estimates (**Figure 2B**), the pattern is less apparent, but can still be observed. For monocular blindness, the majority of tracts revealed a negative effect size (i.e., decreased MD in comparison to controls). In opposition, the majority of tracts in both glaucoma cohorts revealed a positive effect size (i.e., increased MD). This suggests that in monocular blindness a decreased FA in vision tracts is accompanied by an increased FA in non-vision tracts (this could be interpreted as a plasticity and compensatory process). In glaucoma instead no plasticity was observed in non-vision tracts.

### Does glaucoma affect the WMM similarly to monocular blindness?

Next, we wanted to test whether the effect of glaucoma on the WMM is primarily due to reduced visual input. To do so, we set out to determine whether glaucoma and monocular blindness affect the WMM similarly. We remind the reader that monocular blindness was used in our study as a model of decreased visual input. When focussing on FA we found a main effect (F(2,90)=59.29, p<0.0001) indicating a difference between GL1 vs HC1 (g=−0.25 SE=0.04), GL2 vs HC2 (g=−0.28, SE=0.04) and MBL vs HC1 (g=0.28, SE=0.04). Also for MD the main effect was significant (F(2,90)=62.52, p<0.0001), indicating a difference between GL1 vs HC1 (g=0.19 SE=0.04), GL2 vs HC2 (g=−0.08, SE=0.04) and MBL vs HC1 (g=−0.37, SE=0.04). To determine which group comparisons differed from each other, post-hoc comparisons were made using the Tukey HSD test. These tests indicated that the effect size of the glaucoma patients differed from that of MBL group (FA p<0.0001). We repeated the same analysis for our second cohort of glaucoma patients and found that the effect size of the glaucoma patients differed from the MBL group as well (FA p<0.0001). Furthermore we compared effect size for FA and MD between the two glaucoma cohorts and found no difference overall for (FA: p=0.8638 and a small but significant difference for MD: p=0.0043). This suggests that the glaucomatous- and visual-deprived brain respond differently to the loss of visual input.

### Are white matter tract categories similarly affected?

Next, we were interested in establishing whether the WM tract categories were similarly or different in the group comparisons used to estimate g. For FA, a significant main effect of category was found F(6,90)=16.14, p<0.0001, indicating a difference between tract mean g for each WM tract category – early vision (M=−0.65 SE=0.08), ventral stream (M=−0.41, SE=0.06), occipital (M=−0.20, SE=0.07), dorsal (M=−0.01, SE=0.05), vertical (M=−0.0, SE=0.04), callosal (M=−0.02, SE=0.06) and other tracts (M=−0.04, SE=0.04). For MD, a significant main effect of category was found F(6,90)=6.69, p<0.0001, indicating a difference between tract mean g for each WM tract category – early vision (M=0.24 SE=0.08), ventral stream (M=0.10, SE=0.06), occipital (M=−0.12, SE=0.06), dorsal (M=−0.03, SE=0.05), vertical (M=−0.0, SE=0.04), callosal (M=−0.17, SE=0.06) and other tracts (M=−0.15, SE=0.04). These results show that both for FA and MD there was a significant difference in effect size across WM tract categories. This difference in effect size across tract categories suggests that the impact of each disease to tracts beyond the early visual white matter is not uniform.

### Which white matter tract categories are different between group comparisons?

Finally, we were interested in determining which WM tract categories differed between group comparisons. We first plotted the Hedges’ g organized by tract category and group comparison for FA (**Figure 3A**) and MD (**Figure 3B**). We found a significant interaction effect between group comparisons and the tract categories (*F(12,90)=3.23, p*<0.0007). To determine which tract categories differed between group comparisons based on the g estimates of FA, post hoc analysis was performed using Tukey HSD. Our post hoc analysis revealed that both glaucoma group comparisons differed from monocular blindness in tracts of the ventral and dorsal stream and callosal, vertical and other tracts (all *p*<0.0024; **Figure 3A** and **Table 5**). The early vision and occipital WM structures did not reveal differences between any of the group comparisons. None of the categories differed between the two glaucoma group comparisons. The g estimates of MD did not reveal a significant interaction effect between group comparisons and tract categories (**Figure 3B**).

**Figure 3.**
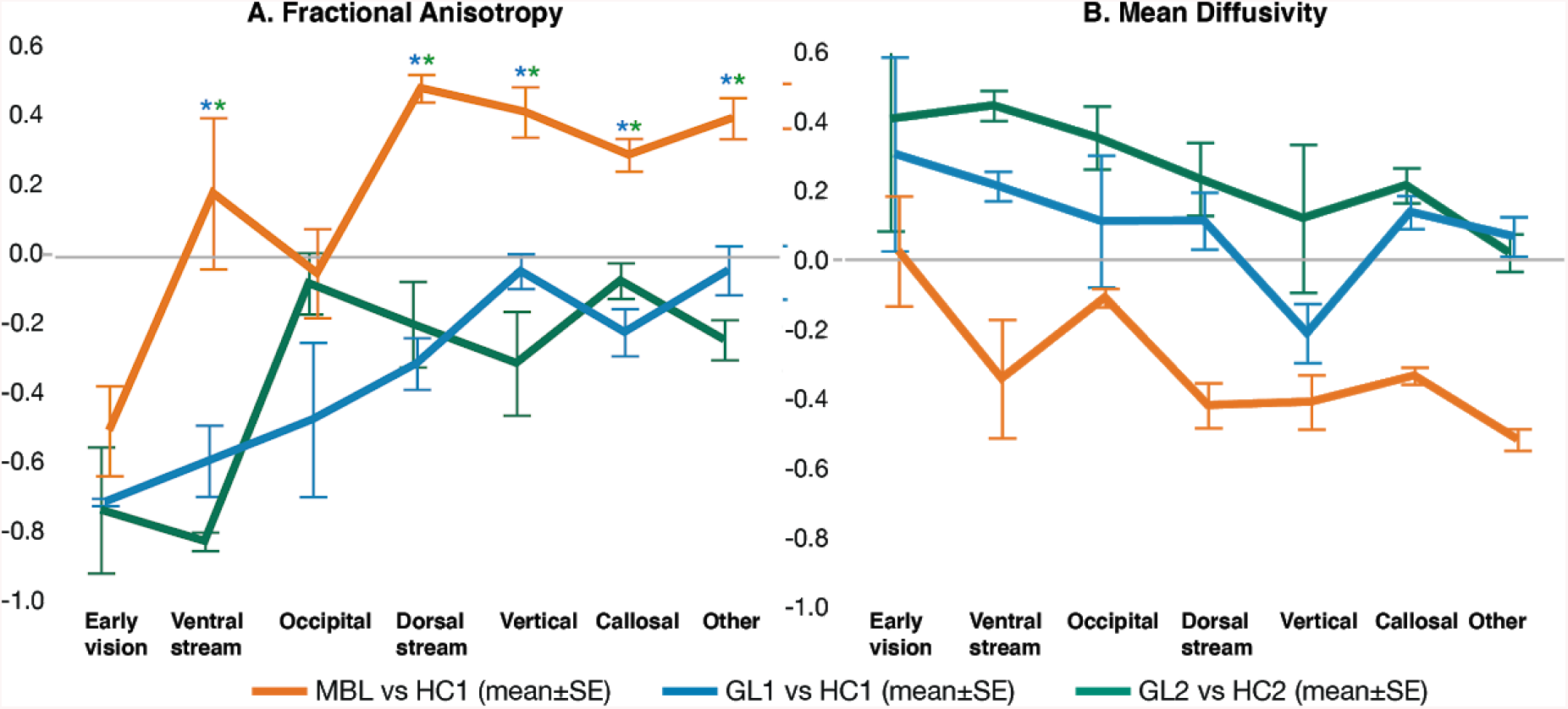
Effect size on the white matter microstructure across tract categories. A. Fractional Anisotropy (FA). B. Mean Diffusivity (MD). The effect of glaucoma and monocular blindness on the WM microstructure compared to healthy controls is visualized for each tract category (abscissa) and participant groups (colors). Positive effects sizes indicate that the microstructural estimate (of either FA or MD) was increased in the clinical group as compared to the healthy controls. Asterisks indicate significance for the comparisons between conditions (colors) with Bonferonni corrected p-values (p < 0.05 / 21 (category*group) = p < 0.0024). The color of the asterix indicates conditions (colors) were different.

**Table 5.** Significant differences between groups per category based on the effect size of fractional anisotropy Results of the post hoc analysis using Tukey HSD to determine which tract categories differed between group comparisons based on the g estimates of fractional anisotropy.

| Category | Group | Group -1 | Diff | SE Diff | Lower CL | Upper CL | p |
| --- | --- | --- | --- | --- | --- | --- | --- |
| **Ventral stream** | MBL vs HC1 | GL2 vs HC2 | 1.0034 | 0.1467 | 0.7119 | 1.2949 | <0.0001 |
| **Ventral stream** | MBL vs HC1 | GL1 vs HC1 | 0.7714 | 0.1467 | 0.4799 | 1.0629 | <0.0001 |
| **Dorsal stream** | MBL vs HC1 | GL2 vs HC2 | 0.6803 | 0.1198 | 0.4423 | 0.9183 | <0.0001 |
| **Dorsal stream** | MBL vs HC1 | GL1 vs HC1 | 0.7936 | 0.1198 | 0.5556 | 1.0316 | <0.0001 |
| **Callosal** | MBL vs HC1 | GL2 vs HC2 | 0.7245 | 0.1467 | 0.4330 | 1.0160 | <0.0001 |
| **Callosal** | MBL vs HC1 | GL1 vs HC1 | 0.4592 | 0.1467 | 0.1677 | 0.7507 | 0.0024 |
| **Vertical** | MBL vs HC1 | GL1 vs HC1 | 0.5113 | 0.1038 | 0.3052 | 0.7174 | <0.0001 |
| **Vertical** | MBL vs HC1 | GL2 vs HC2 | 0.3626 | 0.1038 | 0.1564 | 0.5687 | 0.0007 |
| **Other** | MBL vs HC1 | GL2 vs HC2 | 0.6391 | 0.0928 | 0.4547 | 0.8234 | <0.0001 |
| **Other** | MBL vs HC1 | GL1 vs HC1 | 0.4387 | 0.0928 | 0.2544 | 0.6231 | <0.0001 |

When focussing on FA, Hedges’ g was strongest and negative for the OR (early vision), with a magnitude similar across group comparisons (**Figure 3A**; green, orange and blue). This indicated that FA for the OR was reduced for all patient groups. Hedges g for the rest of the tract categories was strikingly different between group comparisons. Notably, whereas the effect size was primarily positive for the comparison between the monocular blind patients and controls (**Figure 3A** orange), g was primarily negative for the comparisons between the two glaucoma groups and matched controls (**Figure 3A** green and blue). A positive Hedges g for the monocular blind participants can be interpreted as potential indication for WM reorganization processes beyond the visual system as result of prolonged visual input reduction. The opposing pattern of results in the two independent glaucoma group comparisons can be interpreted as an indication of a possible lack of reorganization in the two glaucoma patient groups. The pattern of results for MD is similar –but opposite in the sign of g. Namely, whereas the two glaucoma groups have similar and positive g values in the majority of the tract categories, the monocular blinds participants have negative g for the majority of the tract categories. As in the case of FA also for MD we found no difference between the Hedges g across the three group-comparisons of the early visual WM (i.e., the OR).

## Discussion

This study set out to investigate the effect of glaucoma on the human WM as it is an ongoing debate whether glaucoma is purely an eye disease or whether it affects the whole brain. To address this, we characterized whether the effect of glaucoma on the brain WM is comparable for WM tracts supporting communication within the visual system and for WM tracts serving brain structures beyond the visual system. We used two independent data sets to validate our results.

We used two independent glaucoma datasets and the overall pattern of results was consistent between the two datasets. The effect of glaucoma on WM was strongest in the early visual pathways and it decreased as the tracts of interest were categorized as less involved in serving the visual system. We also used an additional control group of monocular blind individuals. This group was used as a model of reduced visual input to the brain. We compared the pattern of results in the three participant cohorts. We estimated which tracts and categories were most affected in the glaucoma participants and monocular blind patients to test whether any change in the WMM in the glaucoma brain could be explained primarily by a reduction of visual input. We found that whereas the pattern of effects within the two glaucoma patients group was very similar, the effect was strikingly different from that measured in the monocular blind subjects. The results indicated that in monocular blind patients processes of reorganization of the large set of tracts WMM: FA was increased and MD decreased for WMtracts outside of the visual system. This can be cautiously interpreted as plasticity and reorganization following long-term visual deprivation to possibly aid behavior (even though we have no direct evidence that the pattern of alterations in FA and MD relate to behavior). In the two glaucoma groups we found no evidence for a similar reorganization process in the white matter. We found that in both glaucoma groups, FA was reduced and MD increases across all tract categories as compared to controls. Overall, these results show that glaucoma shows a clear difference in the majority of the WM tracts with our model of reduced visual input (the monocular blind group). Below, we will discuss the current findings, the relevance for understanding the aetiology of glaucoma and its clinical relevance.

As it becomes apparent from our results from two independent cohorts − collected at different sites with different parameters − glaucoma consistently shows that WM microstructure is affected strongest in early vision WM structures which then gradually decreases towards cognition structures. The involvement of the early vision structures is consistent with previous studies (C. Boucard et al., 2016; C. C. Boucard et al., 2009; W. W. Chen et al., 2013; Z. Chen et al., 2013; Dai et al., 2013; Garaci et al., 2009; Haykal et al., 2019; Hernowo et al., 2011; C. Li et al., 2012; K. Li et al., 2014; Liu et al., 2012; Murai et al., 2013; Prins et al., 2016; M.-Y. Wang et al., 2013; Yu et al., 2013; Zikou et al., 2012). Our novel findings of a gradually decreasing WM integrity from vision to cognition (**Figure 3**) in glaucoma but not in monocular blindness, indirectly supports the network degeneration hypothesis (NDH). NDH suggests that neurodegenerative diseases primarily spread through distinct brain networks, characteristic of the disease (Tahmasian et al. 2016; Greicius & Kimmel 2012). In light of NDH hypothesis, the RGCs, the visual pathways and the vision-related WM tracts (such as the OR, VOF and ILF) are the primary network for glaucoma and the spread of degeneration among these structures could be part of its clinical manifestation. In regards to glaucoma as a more general neurodegenerative disease, our findings show similarities with involved WM tracts in posterior cortical atrophy (PCA) – a subtype of dementia also referred to as “the visual variant of AD” (Levine et al. 1993; Bokde et al. 2001; Lee & Martin 2004). PCA primarily manifests itself as degeneration of the occipito-temporal (ventral stream) and occipito-parietal (dorsal stream) thereby causing both visuospatial and visuoperceptual problems (Crutch et al. 2012). Furthermore, previous literature has suggested that glaucoma and AD are connected through the same underlying pathologic mechanism (Ghiso et al. 2013, Inoue, Kawaji & Tanihara 2013, Janssen et al. 2013, Sivak 2013). However, the latter notion is still being questioned since conflicting epidemiologic reports have been reported. Some of these epidemiologic studies find an increased prevalence of glaucoma in AD (Bayer, Ferrari & Erb 2002, Chandra, Bharucha & Schoenberg 1986, Helmer et al. 2013, Tamura et al. 2006), while other studies do not (Bach-Holm et al. 2012, Kessing et al. 2007, Ou et al. 2012). Other shared characteristics between AD and glaucoma are the observation of abnormal tau protein in the vitreous jelly of the eyes of glaucoma patients as amyloid beta protein in the retina which are both considered pathologic hallmarks of AD (Criscuolo et al., 2017).

An interesting result to emerge from the data is that glaucomatous WM alterations are significantly different from non-glaucomatous alterations of the WM. This suggests that dissimilarities between glaucoma and decreased visual input alterations likely reflect different underlying mechanisms. In both glaucoma group comparisons, on average, a negative effect size for FA and a positive effect size for MD was observed indicating increased and decreased values in glaucoma patients compared to healthy controls respectively, while the opposite effect is observed for monocular blindness. Glaucoma groups did not differ from each other, despite the fact that glaucoma groups had different pathology (NTG vs monocular POAG). This strengthens the results of comparable patterns of alterations. As little is known about mechanisms that play a role, the discrepancies between glaucoma and monocular blindness groups may favor the hypothesis that WM alterations are not caused by decreased visual input alone, likely reflecting different underlying mechanisms. Although the interpretation of diffusion measures is not straightforward (Beaulieu, 2002; Le Bihan & Van Zijl, 2002), FA is assumed to be highly sensitive for microstructural changes in general, other scalars are more specific for the type of microstructural changes. MD is thought to be sensitive to cellularity, necrosis and edema (Alexander et al., 2007; P. J. Basser et al., 2000; Cornelissen, 2015; Jones et al., 1999). We speculate that on one hand, decreased FA decreases and increased MD in glaucoma may reflect axonal degeneration and demyelination, respectively. On the other ha(Mackey et al., 2012)nd, in monocular blindness the decreased MD and increased FA may reflect WM maturation and high myelination, respectively (Alexander et al., 2007; P. J. Basser et al., 2000; Jones et al., 1999). In support of this latter notion, decreases in MD and increases in FA have been observed after training, supporting the idea that this reflects neuroplasticity or reorganization (Mackey et al., 2012). Our findings corroborate a meta-analysis of 10 dMRI studies examining diffusion values (i.e. FA and MD) in patients with glaucoma, revealing FA decreases and MD increases in the early vision structures. Therefore, we hypothesize that glaucomatous changes are a consequence of a disease progress that involves neurodegeneration, whereas in monocular blindness WM alterations are a consequence of neuroplasticity and reorganization following loss of input.

Our research advances the scientific understanding of glaucoma by using open science methods and data sharing. In addition to improving understanding WM alterations and their potential underlying mechanisms in glaucoma, we have used the open cloud computing platform brainlife.io for secure neuroscience data analysis. The brainlife.io platform allows publishing processed data and associated analyses steps (Apps) integrated into a single record referenced by a digital-object-identifier (DOI) (Avesani et al., 2019). This is a major advancement in supporting reproducibility efforts because it removes the complexities involved with modifying the computational environment for new software, increasing the ability of researchers to easily replicate or add to prior work. All of our data and methods are fully available to the wider scientific community on https://doi.org/10.25663/brainlife.pub.18.

Our research has several clinical implications. First, affected WM integrity shows that damage extends beyond the visual pathways in glaucoma. In addition, we show that monocular glaucoma and monocular blindness affects the WM differently. Determining the full extent of the alternation to the brain WM due to glaucoma could guide the future developments, for example suggesting the span of brain tissues to best target with neuroprotective treatments (Baneke et al., 2020; W. Chen & Wang, 2019; Doozandeh & Yazdani, 2016; Fleischman & Rand Allingham, 2019). For glaucoma, such neuroprotective medication is currently under development (Doozandeh & Yazdani 2016). Neuroprotection may either prevent healthy RGCs from dying or “slow down” the process of already sick RGCs. Combining neuroprotection with IOP reduction - which is clinically proven to prevent RGC loss in glaucoma - may therefore optimize treatment.

As for all dMRI studies, the interpretation of differences between groups in diffusion values is not straightforward (Beaulieu, 2002; Le Bihan & Van Zijl, 2002). Differences may reflect WM abnormalities, but can also be affected by the anatomical features of a tract, such as its curvature, by partial volume effects with neighbouring fiber tracts due to crossing, and by merging or kissing fibers. Fibers originating from the occipital lobe can merge, kiss and cross with fibers from neighbouring tracts (Catani et al., 2003). For example, WM alterations in the OR may consequently have caused (part of) the diffusion changes in neighbouring tracts such as the forcep major as well. However, changes in merging or crossing of fibers could also be part of clinical manifestation itself. Further, for our first cohort (NL), the T1-weighted image had a relatively low contrast ratio between grey- and WM compared to the second cohort (JP). The T1-weighted image is used for segmentation of ROIs and to create a GMWMI mask to constrain tractography. We resolved this by using different WM intensity level settings for each dataset. Although we visually checked all segmentations and masks, a lower contrast ratio between grey- and WM in the first cohort could result in misclassifications in WM segmentation.

In conclusion, glaucoma is associated with widespread WM reduction in fractional anisotropy and increase in mean diffusivity in both early vision as well as non-vision related white matter tracts. Although the early vision and ventral tracts are affected in monocular blindness as well, the WM changes in glaucoma show a different and opposing pattern. Finding such widespread abnormalities in glaucoma suggests the presence of degeneration beyond what can be explained on the basis of decreased visual input. Together with degeneration of the RGCs, the degeneration of visual pathways and ventral WM tracts may be part of the manifestation of glaucoma.

## Materials and methods

### Data sources and App processing records

All data are de-identified and published online using the brainlife.io platform. Data and Apps are interlinked and each dataset is preserved with associated provenance information to allow other researchers to download the minimally processed or processed data and to reuse the Apps on the cloud platform on their own data. Data and Apps associated with the present project are published and archived at https://doi.org/10.25663/brainlife.pub.18.

### Study participants

Data from two sites were included in the current study. Data were collected among the ophthalmologic patient populations of the University Medical Center Groningen (Groningen, the Netherlands (NL)) and Jikei University hospital (Tokyo, Japan (JP)). This resulted in a total of two patient groups and three control groups. The first cohort included 17 NL participants with primary open angle glaucoma (POAG) with a monocular visual field defect (GL1), 20 healthy age-matched NL participants (HC1) and 14 NL age-matched participants with non-glaucomatous monocular blindness (MBL). A second glaucoma cohort was included to validate our results. For the second cohort, we included 30 JP participants with glaucoma with a high prevalence of normal tension glaucoma (NTG) (GL2) and 21 healthy age-matched JP participants (HC2). The first cohort has been previously studied using morphometry of structural MRI data (T1-weighted images) (Prins, 2016), whereas the second cohort have been previously studied using Tract-Based Spatial Statistics (TBSS) (C. Boucard et al., 2016).

The inclusion criteria can be found in **Supplement 1**. Characteristics of the NL participants can be found in **Table 1** and of the JP participants in **Table 2**. This study conformed to the tenets of the Declaration of Helsinki. The study on the NL participants was approved by the medical ethical committee (METC) of the University Medical Center Groningen while that on the JP participants was approved by the METC of the Jikei University School of Medicine. All participants gave their informed written consent prior to participation.

### Clinical data acquisition

*Visual acuity* was measured for all NL participants with a Snellen chart with optimal correction for the viewing distance.

*Visual fields* of the GL1, GL2 and HC2 participants were assessed using a Humphrey Field Analyser (HFA; Carl Zeiss Meditec, Dublin, CA, USA) 30-2 Swedish Interactive Threshold Algorithm SITA) fast. An abnormal visual field was defined as a glaucoma hemifield test ‘outside normal limits’ for at least two consecutive fields; defects had to be compatible with glaucoma and without any other explanation; for a normal visual field the glaucoma hemifield test had to be within normal limits. Visual field loss is reported in terms of mean deviation (VFMD) The Frequency Doubling Technology perimeter (FDT; C20-1 screening mode) was used in the MBL and HC1 participants. An abnormal test result was defined as at least 1 reproducibly abnormal test location at P<0.01. Both eyes of the HC1 participants and the contralateral eye of the MBL participants had to be normal.

*Retinal nerve fiber layer (RNFL)* was assessed for the GL1 patients using laser polarimetry (GDx; Carl Zeiss Meditec, Jena, Germany); outcome measure was the Nerve Fiber Indicator (NFI) (a summary value that indicates the likelihood of glaucomatous RNFL loss). RNFL thickness of the GL2 patients was measured by means of Optical Coherence Tomography (OCT; Stratus OCT 3000; Carl Zeiss Meditec, Dublin, CA, USA).

### Neuroimaging data acquisition

#### Cohort 1 (collection site: Netherlands)

NL participants were imaged using a 3.0 T Philips Intera scanner (Philips, Eindhoven, The Netherlands) . An 8 channel head coil was used. Diffusion-weighted magnetic resonance imaging (dMRI) data were collected with two phase-encoding schemes, i.e anterior-posterior (AP) and posterior-anterior (PA). The following parameters were used for the dMRI pulse sequence using DwiSE technique: 60 diffusion-weighting directions including an additional b0 image, resolution = 128 × 128, APP fat shift = 11.722 pixels, degree of angulation= 0.57 1.405 −15.9, repetition time = 5485 ms, field of view = 240 × 102 × 240, echo time = 79 ms, diffusion b value = 800s/mm2, EPI factor = 45 and voxel dimension = 1.875 × 1.875 × 2.000 mm. The second DW scan has similar parameters as the first, except for: APA fat shift = 2.035 pixels and repetition time = 5516 ms. One T1-weighted (T1w) anatomical image was acquired for each participant using T1W/3D/TFE-2 sequence and following parameters: flip angle = 8 degrees, TR = 8.70ms, TE = 2.98ms, matrix size = 256×256, field of view = 230 × 160 × 180, number of slices =160 slices and voxel size = 1×1#x00D7;1mm3.

#### Cohort 2 (collection site: Japan)

JP participants were imaged using a 3.0 T Siemens MAGNETOM Trio A Tim System scanner (Siemens, Erlangen, Germany) with a 32 matrix head coil. Diffusion-weighted magnetic resonance imaging (dMRI) data were collected. The following parameters were used for the dMRI pulse sequence using single shot echo planar sequence. The scanning parameters were: 64 diffusion-weighting directions including an additional b0 image, TR = 8800 ms, TE = 87 ms, field of view = 210~230, matrix = 140, slice distance factor = 5%, voxel size = 1.5×1.5×1.575mm, b value = 1000 s/mm2 and acceleration factor = 2 (GRAPPA). One T1-weighted (T1w) anatomical image was acquired for each participant using the following T1W 3D MPRAGE sequence using the following parameters: flip angle = 9 degrees, TR = 2300ms, TE = 2.98ms, matrix size = 256×256, field of view = 256, number of slices = 176 slices and voxel size = 1×1#x00D7;1mm3. The brain images were corrected for intensity inhomogeneity using the built-in console.

### Data processing using open cloud computing services on brainlife.io

All data was processed using the reproducible, open cloud computing services available at brainlife.io (Avesani et al., 2019; Stewart et al., 2015; Towns et al., 2014). The workflow used consisted of a series of 20 brainlife.io Apps (**Table 3** in the **Results** section) and is described in detail below. All the Apps used for this study can be accessed at https://doi.org/10.25663/brainlife.pub.18. **Table 3** reports the DOIs to reproducible web services on brainlife.io and the URLs to the source code on GitHub.com.

### Anatomical data (T1w) processing

The T1w anatomical images were preprocessed using the FMRIB Software Library (FSL) function fsl_anat (brainlife.app.273). Using this app, the T1w anatomical images were reoriented and linearly-aligned to the MNI152 1 mm atlas using flirt, and non-linearly aligned to the same atlas using fnirt. Subsequently, the anatomical images were segmented into multiple tissue types (e.g. cortical grey matter, subcortical grey matter, WM, cerebrospinal fluid and pathological tissue) using MRtrix3’s 5ttgen function (R. E. Smith et al., 2012) ([brainlife.app.239](https://github.com/brainlife/app-mrtrix3-5tt)). A mask representing all of the voxels of grey matter-white matter interface (GMWMI) was created and used to seed tractography (see Methods: White matter microstructure modelling). Additionally, the aligned T1w anatomical image was used for segmentation and surface generation using Freesurfer’s recon-all function (Fischl, 2012) (bl.app.0). For the NL dataset, we used a WM intensity level of 105 using the flag -seg-wlo. Next, the bilateral lateral geniculate nuclei were segmented using Freesurfer’s developmental segment_thalamic_nuclei function (Iglesias et al., 2018) (brainlife.app.222). To be able to reconstruct regions of interest (ROIs) for tractography of the OR, population receptive field (pRF) maps were fit to the cortical surfaces generated from Freesurfer and a retinotopy atlas (brainlife.app.187). This app performs a retinotopic mapping in visual areas V1, V2, and V3, as well as higher-order visual areas, from the T1w anatomical images, using an open source library (github.com/noahbenson/neuropythy) (Benson et al., 2012).

### Diffusion data (dMRI) processing

The DWI images were preprocessed using the procedure *dwiprepoc* provided by MRtrix3 (bl.app.68). The app is largely based on the DESIGNER pipeline (Diffusion parameter EStImation with Gibbs and NoisE Removal), developed to identify and minimize common sources of methodological variability. The steps consisted of performing PCA denoising, correction on Gibbs ringing artifact, motion and eddy current correction with FSL, bias correction with ANTs N4 and removal of Rician background noise. For the NL data, AP and PA diffusion scans were combined into a single corrected one by estimating the susceptibility-induced off-resonance field using TOPUP (Jenkinson et al., 2012; S. M. Smith et al., 2004). DWI slices whose average intensity is at least four standard deviations lower than the expected intensity were considered as outliers and replaced with predictions made by the Gaussian Process. The DWI data and gradients were aligned to the ACPC aligned T1-weighted anatomical scan using FSL’s BBR. The final isotropic voxel dimension in mm was set to 1.875 and 1.5 for the NL and JP data, respectively.

### Region-of-interest (ROI) generation

A multiple region of interest (ROIs) approach was used to be able to reconstruct the OR using the roiGenerator app (brainlife.app.223). This app uses AFNI’s 3dROIMaker functionality (Cox, 1996) to generate ROIs in dMRI space by taking the Freesurfer’s parcellation volume, thalamic nuclei segmentation and pRF visual area segmentation, to generate exclusion ROIs, the lateral geniculate nuclei (LGN) and V1, respectively. Exclusions ROIs used from the Freesurfer’s aseg segmentation were as follows: the bilateral cerebral WM, cerebral cortex, lateral ventricle, cerebellum WM and cerebellum cortex.

### White matter microstructure modelling (Tractography)

Probabilistic whole-brain fiber tracking was performed using the MRtrix3 app (Tournier et al., 2012) (bl.app.101) which uses MRtrix3’s Anatomically Constrained Tractography (ACT) algorithm (R. E. Smith et al., 2012). Constrained spherical deconvolution (CSD) was used to estimate local fiber orientation distributions (FOD) (Tournier et al. 2007). CSD estimates were generated with a maximum spherical harmonic order of Lmax = 8. The GMWMI mask was used as a seed mask to constrain fiber tracking. The following parameters were used: a maximum curvature angle of 45° between successive steps, *a step size of 0.2 mm*, a maximum track length of 250 mm and a minimum length of 29 mm. A total of 1,500,000 streamlines per parameter combination was produced.

### White matter microstructure modelling (Tracking the Human Optic RAdiation (THORA))

The OR was reconstructed using the THORA app (brainlife.app.347). This app will generate ORs using MRtrix3’s iFOD2 algorithm with the capability of using the ACT framework and performs backtracking between the bilateral LGN and bilateral V1 as an end ROI. Previously generated exclusion ROIs were used to exclude fibers that are not part of the OR. The following parameters were used: a maximum spherical harmonic order of Lmax = 8, number of repetitions, minimum and maximum length of streamlines of 10 and 180 mm, respectively, a step size of 0.5 mm, a streamline count of 1000, a minimum FOD amplitude of 0.05 as a cutoff and a maximum curvature angle of 35°.

### White matter microstructure modelling (Segmentation)

To examine WM differences in early vision, ventral stream, dorsal stream, occipital, vertical, callosal and other tracts, we identified and segmented a total of 37 WM tracts (see **Table 4 i**n the **Results** section) using the app WM anatomy segmentation (Bullock et al., 2019) (brainlife.app.188). This app Automatically segments a tractogram into the major WM tracts using the Freesurfer 2009 parcellation and the ACT tractography. For an overview of the segmented tracts see **Figure 1** in the **Results** section.

### Remove tract outliers

To remove spurious fibers, tracts were cleaned by removing fibers using brainlife.io wrapper for mbaComputeFibersOutliers algorithm. It takes the tract classification and prunes classified fibers that are unlike other fibers within the same tract (brainlife.app.195). Tracts that deviate more than 3 standard deviations (SDs) from the core of the tract (i.e. maximum distance) as well as fibers that are more than 4 SDs above the mean length of the tract (i.e. maximum length) were removed.

### Tract profile generation

Tract profile generation is performed after the reconstruction of tracts via tractography and subsequent segmentation. For each of the segmented tract, plots of diffusion metrics (i.e. FA, MD, RD, AD) were created, known as Tract Profiles (Yeatman et al., 2012); bl.app.43). The app utilizes the Compute_FA_AlongFGcommand provided by Vistasoft. For each tract the core representation is estimated by applying a gaussian weight to each streamline based on distance away from the “core”. Each tract is resampled into 100 equidistant nodes along their trajectories and the weighted average metric at each node for each participant can be extracted.

### Quality assurance

To check the quality of our analyses, we generated images of each subject of the T1-weighted scan, tissue type masks, DWI scans and DWI scan overlaid on T1-weighted scan. We further plotted the response function, computed SNR and calculated statistics associated with average streamline characteristics of each segmented tract (i.e. count, volume occupied, average length and length distribution). See **Table 3** in the **Results** section for an overview of Apps used.

### Statistical analyses

#### Extraction of white matter properties of tracts

For each WM tract, we computed diffusion along the tract by resampling each tract into 100 equidistant nodes. At each node, fractional anisotropy (FA) and mean diffusivity (MD) were calculated. Individual tracts were excluded from the analysis if they had less than 50 streamlines (see **Table S3** for overview of excluded tracts per group). Additionally, the first and last 10 nodes of each tract were excluded from the analyses since the ends of the tract may contain ambiguous voxels of the grey-white matter junction. The average FA and MD of each tract was calculated.

#### Effect size calculation

To examine the effect of glaucoma and monocular blindness on the WM microstructure compared to healthy controls, we calculated the effect size for both FA and MD separately between groups using the average tract values. We compared participants of GL1 versus HC1 and MBL versus HC1. To validate results, we additionally compared GL2 versus HC2 in our second cohort. The Effect sizes were computed using Hedges’ G for each tract using The Measures of Effect Size (MES) Toolbox (Hentschke & Stüttgen, 2011) (github.com/hhentschke/measures-of-effect-size-toolbox). Hedges’ g was calculated using the following equation: *Hedges’* 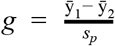 with 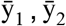 and *s_p_* denoting the mean of sample 1, the mean of sample 2, and the pooled standard deviation, respectively. The equation for the pooled standard deviation is 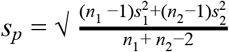 with *s_1_* and *n_1_* denoting the standard deviation and number of observations for sample 1, respectively, and *s_2_* and *s_2_* denoting the standard deviation and number of observations for sample 2, respectively.

Two-Way Factorial ANOVA was conducted independently for FA and MD to compare the main effects of participant group and tract category and the interactions between group and category on the g estimates. The variable group consisted of three different levels (GL1 vs HC1, MBL vs HC1 and GL2 vs HC2) and the variable category consisted of seven levels (early vision, ventral, occipital, dorsal, vertical, callosal and other tracts

##### What are the similarities between glaucomatous WM alterations and those caused by decreased visual input?

To examine whether the effect of glaucomatous alterations is similar to decreased input (as measured by monocular blindness), we investigated the main effect of group for Hedges’ *g* FA and MD, separately, using a Two-Way Factorial ANOVA. The main effect of group consisted of three levels (GL1 versus HC1, MBL versus HC1, GL2 versus HC).

##### Are alterations in the visual WM similar to those in the rest of the WM?

To examine whether alterations in visual WM are similar to the rest of the WM, all WM tracts were categorized based on their involvement in vision. Categorisations were made as follows: 1) early vision tracts are tracts part of the visual pathways (Rokem et al., 2017), 2) ventral stream tracts are associated with ventral stream processing (Goodale & David Milner, 1992), 3) dorsal stream tracts are associated with dorsal stream processing (Goodale & David Milner, 1992), 4) occipital tracts are defined as tracts originating from or projecting to the occipital lobe (Takemura et al., 2017), 5) vertical tracts are defined as vertically-oriented tracts (Bullock et al., 2019), 6) callosal tracts are tracts projecting through the body or genu of the corpus callosum (Fabri et al., 2014)) and 7) other tracts are defined as WM tracts that are not categorized in previous categories. An overview of these categorizations of WM tracts can be found in **Table 4** and **Figure 1** in the **Results** section. Next, the Hedges’ G value for FA and MD of each tract was averaged per tract category to create an average effect size measure of WM property changes in glaucoma groups and non-glaucomatous blindness compared to healthy controls. To examine whether the effect of glaucomatous alterations in visual WM is similar to other WM tracts), we investigated the main effect of WM tract categories for Hedges’ *g* FA and MD, separately, using a Two-Way Factorial ANOVA.

###### Interaction effect group and category

To examine whether WM tract categories are differently affected between groups, we examined the interaction effect between group and categories using Two-Way Factorial ANOVA. If significant at p < 0.05, then Post Hocs using Tukey HSD were performed to examine which categories differed between the groups. If significant at p < 0.05, then posthocs were performed to examine which categories differed within the group. Post Hoc results were Bonferroni corrected for multiple comparisons and significance was set at *p* < 0.05 / 21 (category*group) = *p* < 0.0024.

###### Demographics

Group differences in age and gender were tested using two-sample t-tests and Fisher’s exact tests, respectively.

## Supporting information

S1 Inclusion and exclusion criteria of two datasets

Abbreviations

## Acknowledgements

This research was funded by NSF OAC-1916518, NSF IIS-1912270, NSF IIS-1636893, NSF BCS-1734853, Microsoft Faculty Fellowship to F.P. B.M. was partially supported by NIH NIMH T32-MH103213 to William Hetrick (Indiana University). CCB was supported by a grant from the Japan Society for the Promotion of Science (JSPS) Postdoctoral Fellowship for Overseas Researchers. SH was supported via FWC by research grants from the Stichting MD Fonds, Landelijke Stichting voor Blinden en Slechtzienden (LSBS), and Algemene Nederlandse Vereniging ter Voorkoming van Blindheid (ANVVB) via UitZicht. BCB was supported by NWO VICI grant (No. 453-11-004) awarded to A. Aleman. The funding organizations had no role in the design or conduct of this research. The authors would like to acknowledge the help of Soichi Hayashi and Daniel Bullock for contributing to the development of brainlife.io.

## Author contribution

Study concept and design: SH, FWC, FP. Acquisition of data: DP, CCB, MY, MI, AT. Analysis or interpretation of the data: SH, FWC, FP. Wrote the manuscript: SH, FWC, FP. Critical revision of the manuscript for intellectual content: All authors. Statistical analysis: SH, FP. Technical support: BC, BM, FP. Study supervision: FWC, BCB, NJ, FP

