## Supplementary material for "White matter alterations in glaucoma and vision-deprived brains differ outside the visual system": S1 Inclusion and exclusion criteria of two datasets

**Dutch cohort**

Healthy participants

- Both eyes have an intact visual field. Outcome of Frequency Doubling Technology (C20-1 screenings mode): all test locations are intact (P≥1%).
- A good visual acuity in both eyes. Outcome of the Snellen visual acuity: at least 0.8 (0.1 logMAR or less).

Exclusion:

- Diagnosis with neurological or psychiatric disorders.
- An eye disease.
- Pregnancy.
- Having metal implants or other metal objects in the body.
- Known with claustrophobia .

*Monocular glaucoma participants*

Inclusion

- In one eye a visual field defect due to primary open angle glaucoma, pseudoexfoliation syndrome or pigment dispersion syndrome.
- The contralateral eye has an intact visual field. Outcome of the Humphrey Field Analyzer (30-2 SITA): glaucoma hemifield test 'within normal limits'.
- The contralateral eye has a good visual acuity. Outcome of the Snellen visual acuity: at least 0.8 (0.1 logMAR or less).

Exclusion

- Another eye disease than primary open angle glaucoma, pseudoexfoliation syndrome or pigment dispersion syndrome.
- Pregnancy.
- Having metal implants or other metal objects in the body.
- Known with claustrophobia.
- Diagnosis with neurological or psychiatric disorders.

*Monocular blind participants*

Inclusion

- Unilaterally light-perception negative for at least five years, due to perforation, enucleation, evisceration or ablatio retinae leading to blindness to the affected eye.
- Healthy had to have an intact visual field. Outcome of Frequency Doubling Technology (C20-1 screening mode): all test locations are intact (P≥1%).
- A good visual acuity in the healthy eye. Outcome of the Snellen visual acuity: at least 0.8 (0.1 logMAR or less).

Exclusion

- Existence of monocular vision shorter than 5 years.
- Visus lower than 0.8 in the healthy eye. Outcome of the Snellen visual acuity: at least 0.8 (0.1 logMAR or less).
- Diagnosis with neurological or psychiatric disorders.
- An eye disease.
- Pregnancy.
- Having metal implants or other metal objects in the body.
- Known with claustrophobia .

**Japanese cohort**

*Glaucoma participants*

Inclusion

- Optic nerve head changes consistent with glaucoma assessed with Optical Coherence Tomography (OCT), evaluated by an ophthalmologist
- Visual field defects consistent with glaucoma assessed with a Humphrey Field Analyser (HFA), evaluated by an ophthalmologist

Exclusion criteria

- Diagnosis with neurological or psychiatric disorders.

*Healthy participants*

Exclusion criteria

Diagnosis with neurological or psychiatric disorders.

- An eye disease.
