## Supplemental Table 2 for "White matter alterations in glaucoma and vision-deprived brains differ outside the visual system"

**Table S2** Categorical effect size on the white matter microstructure based on the effect size of fractional anisotropy and mean diffusivity.

| Category | WM Tracts | Fractional Anisotropy | | | Mean Diffusivity | | |
|---|---|---|---|---|---|---|---|
| | | *JP GL vs HC* | *NL GL vs HC* | *NL MBL vs HC* | *JP GL vs HC* | *NL GL vs HC* | *NL MBL vs HC* |
| **Early vision** | OR L | -0.549 | -0.718 | -0.632 | 0.082 | 0.583 | 0.182 |
| | OR R | -0.912 | -0.697 | -0.372 | 0.727 | 0.023 | -0.135 |
| **Ventral stream** | IFOF L | -0.824 | -0.62 | 0.771 | 0.424 | 0.111 | -0.831 |
| | IFOF R | -0.808 | -0.29 | 0.252 | 0.566 | 0.168 | -0.123 |
| | ILF L | -0.761 | -0.72 | -0.111 | 0.361 | 0.28 | -0.332 |
| | ILF R | -0.89 | -0.725 | -0.182 | 0.421 | 0.284 | -0.093 |
| **Dorsal stream** | Cingulum L | 0.279 | -0.129 | 0.659 | -0.19 | -0.143 | -0.201 |
| | Cingulum R | -0.078 | -0.105 | 0.421 | 0.137 | -0.119 | -0.279 |
| | SLF1And2 L | -0.342 | -0.269 | 0.494 | 0.306 | 0.183 | -0.564 |
| | SLF1And2 R | -0.64 | -0.331 | 0.38 | 0.598 | 0.344 | -0.423 |
| | SLF3 L | -0.191 | -0.586 | 0.509 | 0.286 | 0.132 | -0.607 |
| | SLF3 R | -0.197 | -0.428 | 0.452 | 0.25 | 0.269 | -0.456 |
| **Occipital** | Forceps Major | -0.19 | -0.145 | 0.209 | 0.307 | -0.247 | -0.165 |
| | VOF L | -0.139 | -0.368 | -0.172 | 0.526 | 0.172 | -0.078 |
| | VOF R | 0.097 | -0.895 | -0.181 | 0.22 | 0.403 | -0.09 |
| **Vertical** | MDLFang L | -0.069 | 0.015 | 0.448 | 0.303 | 0.263 | -0.22 |
| | MDLFang R | 0.104 | -0.135 | 0.347 | 0.441 | -0.013 | -0.319 |
| | MDLFspl L | 0.033 | 0.006 | 0.166 | 0.115 | -0.115 | -0.403 |
| | MDLFspl R | -0.184 | -0.555 | 0.218 | 0.089 | 0.182 | -0.434 |
| | pArc L | -0.311 | -0.244 | 0.45 | 0.35 | 0.235 | -0.357 |
| | pArc R | -0.167 | -0.163 | 0.403 | 0.214 | 0.253 | -0.372 |
| | TPC L | 0.09 | -0.373 | 0.13 | 0.021 | 0.172 | -0.311 |
| | TPC R | -0.049 | -0.295 | 0.185 | 0.169 | 0.107 | -0.272 |
| **Callosal** | CC AnterioFrontal | -0.207 | 0.061 | 0.478 | -0.009 | -0.399 | -0.411 |
| | CC MiddleFrontal | -0.01 | -0.022 | 0.593 | -0.196 | -0.044 | -0.632 |
| | CC Parietal | -0.72 | -0.027 | 0.28 | 0.748 | -0.096 | -0.285 |
| | Forceps Minor | -0.292 | -0.18 | 0.318 | -0.071 | -0.316 | -0.318 |
| **Other** | Arcuate L | -0.519 | -0.084 | 0.619 | 0.147 | 0.156 | -0.509 |
| | Arcuate R | -0.398 | -0.213 | 0.515 | 0.329 | 0.232 | -0.464 |
| | Aslant L | -0.363 | -0.048 | 0.354 | 0.108 | 0.107 | -0.466 |
| | Aslant R | 0.001 | -0.27 | 0.094 | 0.015 | 0.281 | -0.354 |
| | ATR L | -0.102 | 0.363 | 0.521 | -0.087 | -0.316 | -0.6 |
| | ATR R | 0.025 | 0.095 | 0.603 | -0.233 | -0.114 | -0.5 |
| | CST L | -0.185 | -0.281 | 0.2 | -0.006 | 0.096 | -0.707 |
| | CST R | -0.152 | 0.225 | 0.24 | 0.106 | -0.05 | -0.477 |
| | Uncinate L | -0.362 | -0.248 | 0.292 | -0.223 | 0.064 | -0.502 |
| | Unicinate R | -0.342 | 0.068 | 0.556 | 0.027 | 0.199 | -0.619 |

The effect of glaucoma and monocular blindness on the white matter microstructure compared to healthy controls per category and group. Positive effects sizes indicate that the value of the clinical group (either FA or MD) was increased compared to the healthy controls.
