## Supplemental Table 3 for "White matter alterations in glaucoma and vision-deprived brains differ outside the visual system"

**Table S3 Number of subjects with excluded white matter tract**

| Tract | *GL_JP* | *HC_JP* | *GL_NL* | *HC_NL* | *MBL_NL* |
|---|---|---|---|---|---|
| **L OR** | 5 | 3 | 2 | 1 | 2 |
| **R OR** | 3 | 3 | 3 | 1 | 2 |
| **Forceps Minor** | 0 | 0 | 0 | 2 | 0 |
| **Forceps Major** | 0 | 0 | 0 | 2 | 0 |
| **Parietal CC** | 1 | 0 | 1 | 2 | 0 |
| **Middle Frontal CC** | 0 | 0 | 0 | 2 | 0 |
| **Anterior Frontal CC** | 0 | 0 | 0 | 2 | 0 |
| **L Cingulum** | 3 | 0 | 1 | 4 | 2 |
| **R Cingulum** | 0 | 0 | 1 | 3 | 0 |
| **L Uncinate** | 0 | 0 | 1 | 2 | 2 |
| **R Uncinate** | 0 | 0 | 0 | 2 | 0 |
| **L IFOF** | 5 | 2 | 6 | 8 | 6 |
| **R IFOF** | 4 | 1 | 0 | 3 | 1 |
| **L Arcuate** | 1 | 0 | 0 | 2 | 0 |
| **R Arcuate** | 3 | 0 | 0 | 2 | 0 |
| **L SLF 1/2** | 0 | 0 | 1 | 2 | 0 |
| **R SLF 1/2** | 0 | 0 | 0 | 2 | 0 |
| **L SLF 3** | 0 | 0 | 0 | 2 | 0 |
| **R SLF 3** | 0 | 0 | 0 | 2 | 0 |
| **L Fronto Thalamic** | 1 | 0 | 0 | 2 | 0 |
| **R Fronto Thalamic** | 1 | 0 | 0 | 2 | 0 |
| **L Aslant** | 1 | 0 | 0 | 2 | 0 |
| **R Aslant** | 1 | 0 | 0 | 3 | 0 |
| **L ILF** | 0 | 0 | 0 | 2 | 0 |
| **R ILF** | 0 | 0 | 0 | 2 | 0 |
| **L MDLF Ang** | 1 | 0 | 0 | 2 | 0 |
| **R MDLF Ang** | 1 | 1 | 0 | 2 | 0 |
| **L MDLF SPL** | 5 | 3 | 4 | 2 | 1 |
| **R MDLF SPL** | 3 | 4 | 3 | 3 | 2 |
| **L pArc** | 0 | 0 | 0 | 2 | 0 |
| **R pArc** | 1 | 0 | 0 | 2 | 0 |
| **L TPC** | 2 | 0 | 0 | 2 | 0 |
| **R TPC** | 2 | 0 | 0 | 2 | 0 |
| **L VOF** | 0 | 1 | 0 | 2 | 0 |
| **R VOF** | 0 | 0 | 0 | 2 | 0 |
| **L CST** | 1 | 0 | 0 | 2 | 0 |
| **R CST** | 1 | 0 | 0 | 2 | 0 |

Number of participants with specific tracts excluded in each dataset. Tracts were excluded if they had less than 50 streamlines.
