## Supplemental Table 1 for "White matter alterations in glaucoma and vision-deprived brains differ outside the visual system"

**Table S1** Average effect sizes on the white matter microstructure for fractional anisotropy and mean diffusivity measured across all 37 WM tracts.

| | **Fractional Anisotropy** | | **Mean Diffusivity** | |
|---|---|---|---|---|
| *Comparison* | *Mean (g)* | *Std Error* | *Mean (g)* | *Std Error* |
| **GL1 vs HC1** | -0.25 | 0.04 | 0.09 | 0.04 |
| **GL2 vs HC2** | -0.28 | 0.04 | 0.20 | 0.04 |
| **MBL vs HC1** | 0.28 | 0.04 | -0.37 | 0.04 |

Average effect sizes for each group comparison were calculated based on all white matter tracts. Between group differences were determined using Two-Way (Factorial) ANOVA was used independently for the two glaucoma datasets and respective control groups. The average effect size per group comparison revealed that the effect of glaucoma and monocular blindness on the WM microstructure differed significantly between groups (both FA and MD $F<0.0001$).
