## Supplementary material for "White matter alterations in glaucoma and vision-deprived brains differ outside the visual system": Abbreviations

**List of Abbreviations**

| | |
|---|---|
| AD | Axial Diffusivity |
| Ang | Angular |
| Arc | Arcuate Fasciculus |
| ATR | Anterior Thalamic Radiation |
| CC | Corpus Callosum |
| Cing | Cingulum Cingulate |
| CST | Corticospinal Tract |
| DWI | Diffusion-weighted Magnetic Resonance Imaging |
| FA | Fractional Anisotropy |
| FAT | Frontal Aslant Tract |
| FMaj | Forceps Major |
| FMin | Forceps Minor |
| GL | Glaucoma |
| GMWMI | Grey Matter White Matter Interface |
| HC | Healthy Control |
| IFOF | Inferior Frontal Longitudinal Fasciculus |
| ILF | Inferior Longitudinal Fasciculus |
| JP | Japanese |
| L | Left |
| MBL | Monocular Blindness |
| MD | Mean Diffusivity |
| MLF | Medial Longitudinal Fasciculus |
| MRI | magnetic resonance imaging |
| NL | Netherlands |
| NTG | Normal Tension Glaucoma |
| OR | Optic Radiation |
| pArc | Parietal Arcuate Fasciculus |
| R | Right |

| | |
|---|---|
| RD | Radial Diffusivity |
| SLF | Superior Longitudinal Fasciculus |
| SPL | Superior Parietal Lobule |
| TPC | Temporal Parietal Connection |
| UF | Uncinate Fasciculus |
| VOF | Ventral Occipital Fasciculus |
| WM | White Matter |
